# Metagenomic and Metabolomic Analyses Reveal Synergistic Effects of Fecal Microbiota Transplantation and Anti-PD-1 Therapy on Treating Colorectal Cancer

**DOI:** 10.1101/2022.02.13.480233

**Authors:** Jiayuan Huang, Xing Zheng, Wanying Kang, Huaijie Hao, Yudan Mao, Hua Zhang, Yuan Chen, Yan Tan, Yulong He, Wenjing Zhao, Yiming Yin

## Abstract

Anti-PD-1 immunotherapy has saved numerous lives of cancer patients; however, it only exerts efficacy in 10-15% of patients with colorectal cancer. Fecal microbiota transplantation (FMT) is a potential approach to improving the efficacy of anti-PD-1 therapy, whereas the detailed mechanisms and the applicability of this combination therapy remain unclear. In this study, we evaluated the synergistic effect of FMT with anti-PD-1 in curing colorectal tumor-bearing mice using a multi-omics approach. Mice treated with the combination therapy showed superior survival rate and tumor control, compared to the mice received anti-PD-1 therapy or FMT alone. Metagenomic analysis showed that composition of gut microbiota in tumor-bearing mice treated with anti-PD-1 therapy was remarkably altered through receiving FMT. Particularly, *Bacteroides* genus, including FMT-increased *B. thetaiotaomicron, B. fragilis*, and FMT-decreased *B. ovatus* might contribute to the enhanced efficacy of anti-PD-1 therapy. Furthermore, metabolomic analysis upon mouse plasma revealed several potential metabolites that upregulated after FMT, including punicic acid and aspirin, might promote the response to anti-PD-1 therapy via their immunomodulatory functions. This work broadens our understanding of the mechanism by which FMT improves the efficacy of anti-PD-1 therapy, which may contribute to the development of novel microbiota-based anti-cancer therapies.

## 1. Introduction

The application of immune checkpoint inhibitors (ICIs) has led to remarkable advances in the treatment of a wide range of cancers, including melanoma, non-small-cell lung cancer (NSCLC), gastric cancer, and breast cancer (1). Antibodies targeting the programmed cell death protein 1 (PD-1) are the most widely used ICIs, which work by blocking the binding between PD-1 receptor of T cells and PD-L1 ligand of tumor cells, and restoring the function of T cells that recognizes and eliminates tumor cells (2). ICI therapy has saved numerous lives since its approval in 2014 and could maintain long-term disease control in ICI responders. However, in terms of curing colorectal cancer (CRC), the majority of patients would present non-response to anti-PD-1 treatment due to the insufficient tumor-infiltrating lymphocytes (TILs) in the tumor microenvironment (TME) (3, 4). Only approximately 10% of patients with CRC, which are mismatch repair deficient (dMMR) or microsatellite instability high (MSI-H) subtypes, could benefit from anti-PD-1 therapy (5, 6). Therefore, it is important to develop novel strategies to optimize our current ICI therapy.

Human intestine harbors more than 10^13^ microorganisms, which play a key role in mediating human health and disease via shaping systematic and local immune functions (7). Since 2015, multiple studies have elucidated that the composition of gut microbiota was associated with the efficacy of anti-PD-1 therapy (8, 9). Notably, three groups (10-12) reported their work in 2018 observing highly diversified bacterial features (i.e. high abundance of *Akkermansia, Ruminococcus*, and *Bifidobacterium*) were individually related to the favorable clinical outcomes. The mechanisms by which gut microbiota improves anti-PD-1 efficacy involve the increased abundance of beneficial bacteria, enhancement of dendritic cell (DC) maturation, increased activity of anti-tumor CD8^+^ T cells, and the promotion of T cell tumor infiltration (13). These findings suggest the potential approach to enhancing the effect of immunotherapy via regulating gut microbes (14).

Fecal microbiota transplantation (FMT) is a biomedical technology of transplanting functional microbiota into patients, to cure diseases via rebuilding the intestinal flora with normal functions (12). FMT has been employed clinically as a main or adjunctive approach in treating a number of diseases, including *Clostridium difficile* infection, inflammatory bowel diseases, and irritable bowel syndrome (15). In 2021, two independent clinical studies demonstrated that FMT could promote the efficacy of anti-PD-1 therapy in 3/10 and 6/15 patients with PD-1-refractory melanoma, respectively (16, 17). Genes associated with peptides presentation by antigen-presenting cells (APCs) through MHC class I and IL-1 mediated signal transduction were upregulated in melanoma patients after FMT treatment (16). Another study demonstrated that patients with epithelial tumors who responded to the combination treatment of FMT and ICI exerted increased compositions of CD8^+^ T cells, T-helper type I cells, and APCs in the tumor microenvironment, while a reduction of myeloid-derived suppressor cells infiltration was observed (10). Animal experiments elucidated that fecal transplantation into mouse models for lung cancer led to superior tumor suppression (18). However, the detailed mechanism and the applicability of this combination therapy in other cancer types require to be further illustrated.

In this study, we evaluated the antitumor efficacy of FMT in combination with anti-PD-1 immunotherapy using CRC tumor-bearing mouse models and investigated the underlying mechanisms through multi-omics approaches. Our results provide a mechanistic basis of the synergistic effects of FMT and anti-PD-1 therapy on treating colorectal cancer, which will expand our knowledge on the mechanism of immunotherapy and assist with the development of novel anticancer therapy through modulating microbiota.

## 2. Methods

### 2.1. Animals

All animal experiments were conducted at Crown Biosciences Co. Ltd. (Taicang, China) and approved by its Institutional Animal Care and Use Committee (approval number: E4756-B1901). Female BALB/c mice were purchased from Shanghai Lingchang Biological Technology Co. Ltd. (animal certificate number: 20180003003129). All mice were housed under specific-pathogen-free conditions with ingested pellet food (radio-sterilized with cobalt 60) and autoclaved water provided ad libitum.

### 2.2. FMT production

Stool samples from healthy human donors were collected using sterile boxes and processed within 2 h, as previously described (19). In a sterile anaerobic environment, the samples were thoroughly mixed with sterile normal saline (mass: volume = 1:5). Subsequently, filter bags with apertures of 1 mm, 0.25 mm, and 0.05 mm were used to remove solid particles and impurities in the stool samples. The filtered liquid was centrifuged at 5500 g at 4 °C for 5 min, and the precipitation was collected. Bacterial viable counting was conducted via flow cytometry and anaerobic plate counting. The bacterial solution was adjusted to 0.83×10 ^11^ colony forming units per mL (CFU/mL), and mixed with autoclaved glycerol, frozen at −80°C until next use.

### 2.3. Cell culture

CT26 mouse colon carcinoma cell lines were obtained from the Shanghai Institute of Life Sciences (CAT#: TCM37). Cells were cultured in RPMI 1640 culture medium (Gibco) supplemented with 10% fetal bovine serum (FBS) (Excell) and were cultured in a humidified incubator at 37°C, 5% CO2. CT26 cells at the exponential growth stage were suspended in PBS for subcutaneous tumor inoculation in mice.

### 2.4. Tumor-bearing mouse model

Mice (7-8 weeks old) were inoculated with 5×10^5^ CT26 cells per mouse by subcutaneous injection at Day 0 (**Fig. 1A**). A total of 40 mice were randomly divided into four groups: Saline plus Rat IgG2a (designated as Control), FMT plus Rat IgG2a (FMT), Saline plus PD-1 antibody (aPD-1), and FMT in combination with PD-1 antibody (Combo). Sterile normal saline (200 μL per dose) or FMT (5×10^9^ CFU/mouse) was administered by oral gavage on Days 9, 12, 15, and 18; Rat IgG2a (200 μg/mouse, Lenico) and PD-1 antibody (200 μg/mouse, RMP1-14, Lenico) was given by intraperitoneal injection on Days 8, 11, 14, and 17. On Day 24, the endpoint of the experiment, feces, blood, and tumors of tumor-bearing mice were collected, and tumor volume was determined as length × width^2^ × 0.5. Survival rate was defined as the percentage of mice with a tumor volume of less than 2,000 mm^3^ in each group.

**Figure 1.**
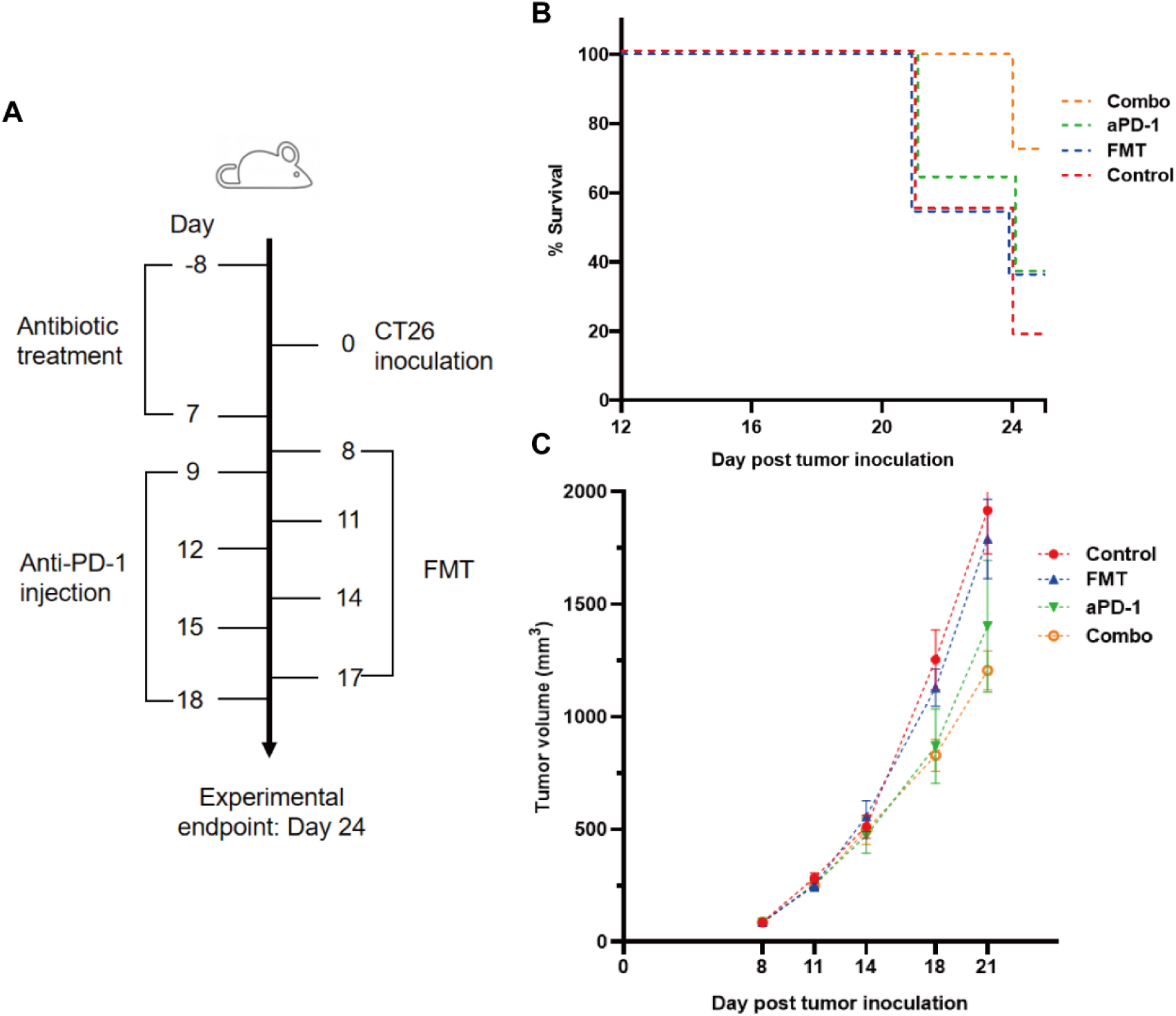
FMT and PD-1 antibody exerted synergistic anti-tumor effect in CT26 tumor-bearing mice. (A) Schematic diagram of this study. (B) Survival curve of the CT26 tumor-bearing mice treated with FMT, aPD-1 or the combination. (C) Tumor growth curves of the CT26 tumor-bearing mice treated with FMT, aPD-1 or the combination. Data are represented as mean ± SD (n = 10).

### 2.5. Antibiotic treatment

From eight days before the tumor inoculation (Day -8) to Day -4, antibiotics were added to the drinking water in proportion, including ampicillin 1 mg/ml, neomycin 1 mg/mL, metronidazole 1 mg/mL, vancomycin 0.5 mg/mL. From Day -3 to Day 7, ampicillin 1 mg/mL was added to the drinking water, and the mixture of metronidazole 10 mg/mL, neomycin 10 mg/mL, vancomycin 5 mg/mL, and amphotericin B 0.1 mg/mL was orally gavaged into each mouse twice a day, 200 μL each time.

### 2.6. Fecal DNA extraction and metagenomic analysis

Total genomic DNA of mouse fecal samples was extracted using QIAamp PowerFecal Pro DNA Kit (Qiagen, CAT#: 51804), according to the manufacturer’s instructions. The concentration was detected by Qubit and the integrity of DNA bands was detected by agarose gel electrophoresis. Library construction and sequencing (Illumina NovaSeq 6000 platform) were performed at Novogene. Following data analyses were performed using Kneaddata, MetaPhlAn and HUMAnN softwares (20).

### 2.7. Metabolomic analysis

Mice blood samples were mixed with ice-cold methanol (3:1, v:v), and centrifuged with 12,000 rpm at 4 °C for 10 min. The supernatant was collected and centrifuged at 12,000 pm at 4 °C for 5 min. The sample extractions were analyzed using an LC-ESI-MS/MS system (UPLC, Shim-pack UFLC Shimadzu CBM A system; MS, QTRAP® System). Chromatographic separation was carried out on a Waters ACQUITY UPLC HSS T3 C18 (1.8 µm, 2.1 mm*100 mm) column. Subsequently, the mass spectrometry separation was carried out using electrospray ionization (ESI) in the positive and negative mode (21). Following data analysis was performed using MetaboAnalyst (22).

### 2.8. Statistical analysis

Statistical analyses were performed using R programming (version 4.0.3) and GraphPad Prism (version 8.0.2). Linear discriminant analysis effect size (LEfSe) was applied to identify differential species based on relative abundance using the Galaxy platform (http://huttenhower.sph.harvard.edu/galaxy). One-way analysis of variance (ANOVA) was performed to illustrate differential bacterial species and serum metabolites among multiple groups. Spearman’s correlation analysis was used to illustrate the relationship between bacterial species and metabolites.

## 3. Results

### 3.1. FMT improved the efficacy of aPD-1 in tumor-bearing mouse model

We evaluated tumor volume and survival rate in CT26 tumor-bearing mice treated with FMT or aPD-1 either alone or in combination (**Fig. 1A**). The Combo group showed the highest animal survival rate (70% vs. 10%, 30%, and 30% in control, FMT, and aPD-1 groups, respectively) on Day 24 after tumor incubation (**Fig. 1B**). Consistently, the Combo group exhibited the most significant tumor growth suppression (47.78% vs 16.18% and 36.74% in FMT, and aPD-1 groups) compared with the Control group, respectively (**Fig. 1C**). These results showed that the combination therapy had a superior effect than either monotherapy alone in treating CT26-bearing mice. Notably, in the FMT monotherapy group, the tumor growth suppression showed no significant difference with that of Control group (p=0.845), indicating FMT exerted its therapeutic effects via a synergistic function with aPD-1, rather than repressing tumor alone.

### 3.2. FMT altered the composition of gut microbiota in tumor-bearing mice treated with aPD-1

To investigate whether FMT improved the effects of aPD-1 by refining the gut microbiome, we next performed metagenomic analysis to examine FMT-induced changes of gut microbial composition and gene function. The PCA plot showed an obvious group-based clustering pattern among groups with or without FMT treatment, indicating that FMT significantly changed the composition of gut microbiota, while the change caused by aPD-1 was less remarkable (**Fig. 2A**). FMT contributed to, at the family level, the decrease of the relative abundance of Bifidobacteriaceae, Porphyromonadaceae, Verrucomicrobiaceae, and the increase of Desulfovibrionaceae and Bacteroidaceae (**Fig. 2B)**.

**Figure 2.**
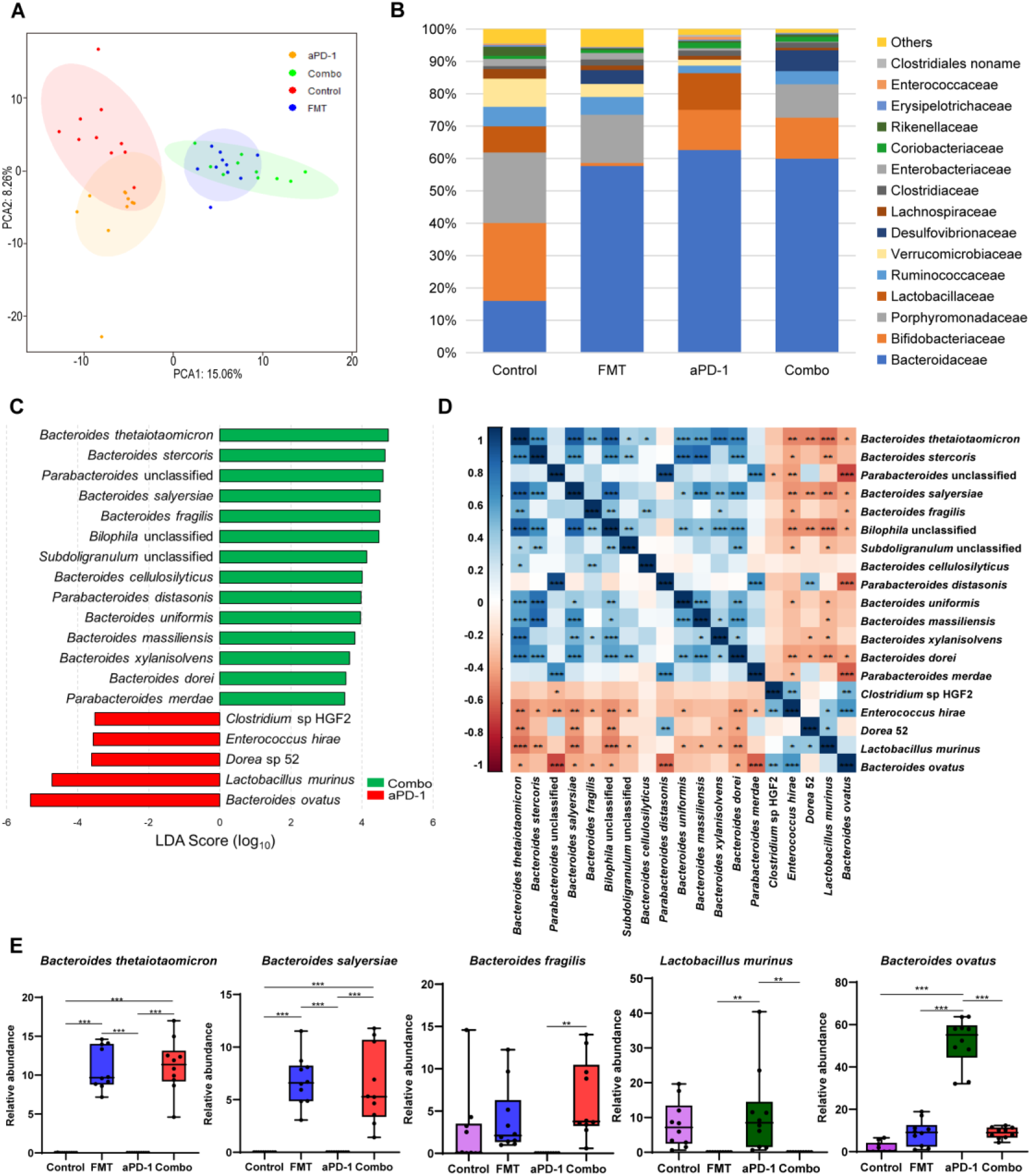
FMT altered the composition of gut microbiota in CT-26 tumor-bearing mice receiving anti-PD-1 therapy. (A) Principal components analysis (PCA) plot of the gut microbiota from mice. (B) Relative abundance of top 15 bacterial families in different groups. (C) LEfSe analysis showing differentially abundant bacterial species between FMT and Combo groups. (D) Heatmap showing the correlations of species significantly different between FMT and Combo groups. (E) Abundance of specific species in different groups. Data are represented as mean ± SD. *, p<0.05; **, p<0.01; ***, p<0.001.

Nineteen significant differentially abundant species between the Combo group and aPD-1 group were identified. The relative abundance of multiple *Bacteroides* species (B. *thetaiotaomicron, B. stercoris, B. salyersiae, B. fragilis, B. cellulosilyticus, B. uniformis*, and *B. massiliensis*) and *Parabacteroides* species (*P. distasonis* and *P*. unclassified) were significantly increased in the mice treated with the combination of FMT and aPD-1, compared to those treated with aPD-1 alone. Also, decreases of the abundance of *Clostridium* sp HGF2, *Enterococcus hirae, Dorea* 52, *Lactobacillus murinus*, and *Bacteroides ovatus* were observed (**Fig. 2C, E, Fig. S1A, B**). In addition, we observed the abundance of specific bacteria, including *Alistipes indistinctus, Faecalibacterium prausnitzii, Bacteroides vulgatus*, and *Oscillibacter* unclassified were enriched, while *Bifidobacterium pseudolongum* were decreased by FMT treatment(p<0.05), and opposite trends were observed in aPD-1 group (**Fig. S1B**).

The abundance of the aforementioned *Bacteroides* species showed a strong positive correlation with each other, as well as a negative correlation with *Enterococcus hirae, Dorea* 52, and *Lactobacillus murinus* (**Fig. 2D**). Interestingly, the abundance of *Bacteroides ovatus* correlated negatively with the abundance of most of the FMT-upregulated species (**Fig. 2D**).

### 3.3. FMT upregulated microbial biosynthetic pathways of nucleotides and amino acids

Other than microbial composition, microbial gene functional changes upon treatments were evaluated. Compared to those of aPD-1 group, 27 differently abundant pathways out of 491 were identified in the combination group (|log2FC|>1, p-adjusted<0.05), indicating the potential microbial contribution towards better anti-PD-1 efficacy induced by FMT (**Fig. 3A**). We observed that the anabolic pathways of several amino acids, including ornithine, histidine, lysine, citrulline, and isoleucine were significantly enriched by FMT treatment. And the pathways of nucleotides de novo biosynthesis, including pyrimidine deoxyribonucleotides, guanosine nucleotides, and adenosine nucleotides were significantly up-regulated in FMT and Combo group. Notably, the pathways of methionine and S-adenosyl-L-methionine (SAM) biosynthesis were significantly decreased, and pathways of S-adenosyl-L-methionine cycle I was increased by FMT treatment. Moreover, the pathways of coenzyme A biosynthesis I, O-antigen building blocks biosynthesis, and heme biosynthesis II were enriched in the aPD-1 group, while down-regulated in the Combo group. Furthermore, the pathway of biotin biosynthesis was significantly up-regulated by FMT treatment (**Fig. 3B, Fig. S2**).

**Figure 3.**
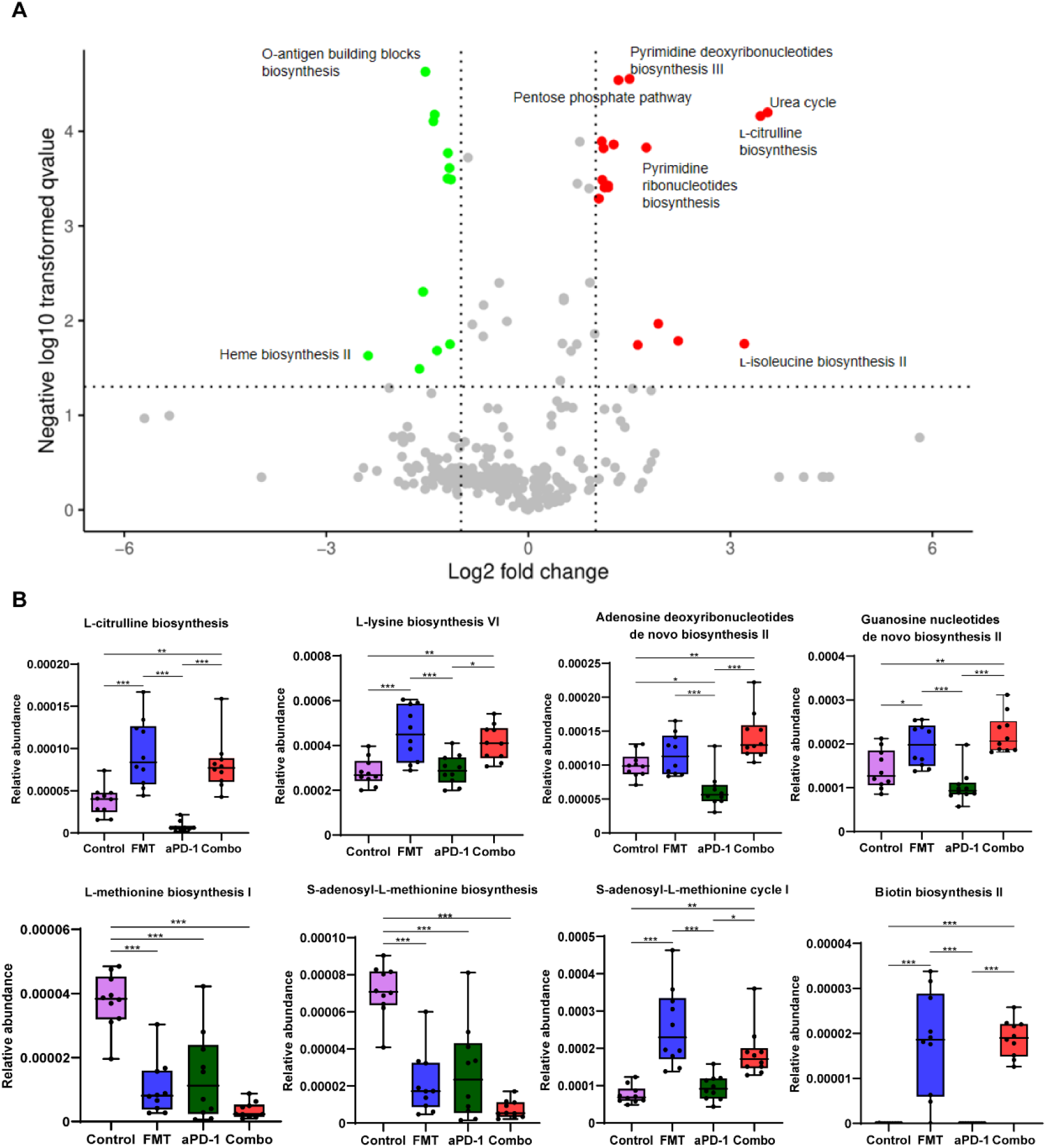
The effect of FMT and PD-1 antibody administration on gut metagenomic gene pathways. (A) Volcano plot showing differentially expressed microbial gene pathways between Combo and aPD-1 groups. (B) Abundance of specific gene pathways in different groups. Data are represented as mean ± SD. *, p<0.05; **, p<0.01; ***, p<0.001.

### 3.4. FMT and aPD-1 synergistically remodeled mouse plasma metabolome

Metabolomic analyses were performed to examine the systemic change caused by FMT in tumor-bearing mice. Among a total number of 369 metabolites detected, the abundance of 8, 9, 34 metabolites were altered following aPD-1, FMT, and Combo treatment, respectively, suggesting the synergistic effect of the combination treatment (**Fig. 4A, D**). Moreover, P-hydroxyphenyl acetic acid, hyodeoxycholic acid, and mandelic acid were decreased in FMT and Combo groups (p-adjusted < 0.05) but not in the aPD-1 group. These metabolites may partially explain the synergistic effect of FMT and aPD-1 (**Fig. 4A**).

**Figure 4.**
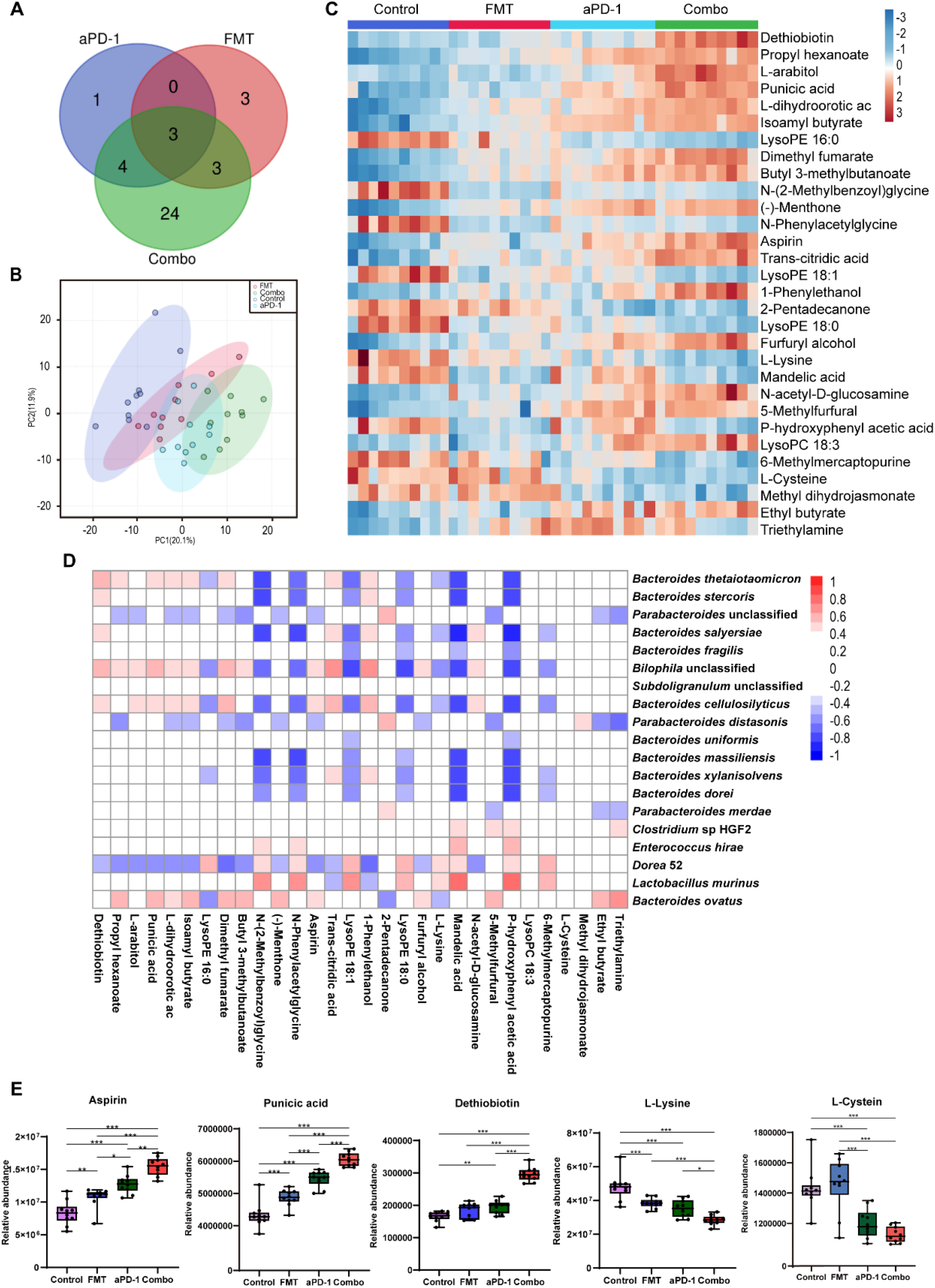
FMT altered plasma metabolites in CT-26 tumor-bearing mice receiving anti-PD-1 therapy. (A) Venn diagrams showing number of significantly changed metabolites in each group after treatment. (B) PCA plot of metabolomic results. (C) Heatmap of differentially abundant metabolites using one-way analysis of variance. (D) The correlations between metabolites and microorganism. (E) Abundance of specific metabolites in different groups. Data are represented as mean ± SD. *, p<0.05; **, p<0.01; ***, p<0.001.

Top 30 most differentially abundant metabolites among the four groups were identified via ANOVA analysis (**Fig. 4C**). Compared with the PD-1 group, dethiobiotin, punicic acid, aspirin, L-arabitol, N-acetyl-D-glucosamine, L-dihydroorotic acid, dimethyl fumarate, trans-citridic acid, 1-Phenylethanol were significantly increased in the Combo group (p<0.01). While lysoPE (16:0), triethylamine, glycine, L-lysine, mandelic acid, L-glutamic acid, L-phenylalanine were significantly decreased (p<0.01) (**Fig. 4C, E, Fig. S3**). The results indicated that combination treat of FMT and aPD-1 significantly altered plasma metabolic profiles. Furthermore, amino acids, including N-(2-Methylbenzoyl) glycine, N-phenyl acetyl glycine, glycine, L-proline, L-cysteine, L-serine and L-lysine were significantly down-regulated in the Combo group (p<0.05). Notably, the abundance of dethiobiotin, propyl hexanoate, and N-acetyl-D-glucosamine were significantly up-regulated in the Combo group (**Fig. 4C, E, Fig. S3**).

To better understand the involvement of specific bacteria species in the alteration of host metabolism, correlation between plasma metabolites and the abundance of specific bacteria species were investigated. High abundance species of *Bacteroides*, such as *B. thetaiotaomicron, B. stercoris, B. salyersiae, B. cellulosilyticus*, were positively correlated with the low abundance of lysoPE (18:0), lysoPE (18:1), N-phenyl acetyl glycine, N-(2-Methylbenzoyl) glycine in plasma, and opposite trends were observed in *B. ovatus* and *Lactobacillus murinus* (**Fig. 4B**). This result suggests a potential link among commensal microorganisms, differentially abundant metabolites, and treatment outcomes of anti-PD-1 therapy efficacy.

## 4. Discussion

Microbiota transplantation of the feces from patients who responded to ICIs combined with ICIs exerts as a promising approach to treating melanoma (17). Yet, the detailed mechanisms and the applicability of this therapy are required to be further evaluated in multiple cancer types, such as colorectal cancer and lung cancer. Moreover, FMT using feces of cancer patients might carry safety risks such as detrimental pathogens or pathobionts; therefore, it’s also necessary to examine the effect of FMT using feces from healthy donors. In this study, our multi-omics investigation shows the synergistic effects of FMT using feces from healthy screened donors and anti-PD-1 therapy, in the treatment of mice bearing colorectal tumor.

A wide range of commensal bacterial species have been reported to enhance the efficacy of ICIs, including *B. thetaiotaomicron* (23), *B. fragilis* (24), *B. cellulosilyticus* (25), *Parabacteroides distasonis* (26), *B. salyersiae* (27), and *B. uniformis* (13). In this study, our metagenomic analysis showed that FMT significantly upregulated the abundance of those aforementioned species, particularly those species from *Bacteroides* genus (**Fig. 2C, E**). The reshaped microbiota caused by FMT might be associated with the refinement of tumor immune microenvironment (TIME) (28). Previous literature shows that *B. thetaiotaomicron*, which is most significantly upregulated by FMT in our data, has been reported to induce immune responses in dendritic cells (e.g. the expression of IL-10) and mediate intestinal homeostasis (29). *B. thetaiotaomicron* is also able to inhibit the growth of CRC cells via its metabolite propionate (23). Another *Bacteroides* species *B. fragilis* is associated with the favorable clinical outcome of CTLA-4 inhibitors (24) via inducing regulatory T cells to secrete IL-10 through the immunomodulatory molecule polysaccharide A (PSA) of *B. fragilis* (30). Additional immunomodulatory function of *B. fragilis* includes producing unique alpha-galactose ceramides (BfaGC) and subsequently activating NKT cells (e.g. upregulating IL-2 expression) (31). More recently, *B. cellulosilyticus* has been reported to be enriched in humanized microbiome mouse model of glioma and is a potential contributor to the enhanced efficacy of anti-PD-1 therapy (25). *B. cellulosilyticus* might modulate host immunity via its specific zwitterionic capsular polysaccharides (ZPSs) which can activate IL-10^+^ regulatory T cells to secrete IL-10 (25). Notably, the abundance of upregulated *Bacteroides* species showed a strong positive correlation with each other (**Fig. 2D**), suggesting their potential symbiotic link. Furthermore, several bacterial species which showed an up-regulation in the Combo group, *Bilophila wadsworthia* and *Lachnospiraceae bacterium* have not been reported previously. Their roles in anti-PD-1 treatment would be very interesting to investigate.

The abundance of two potentially detrimental species, *B. ovatus* and *Lactobacillus murinus*, were significantly decreased by FMT (**Fig. 2C, E**). It was previously reported that the abundance of *B. ovatus* was associated with shorter progression-free survival (PFS) in melanoma patients receiving immunotherapy (32). *B. ovatus* might affect host immunity via inducing IgA and other approaches (33). In addition, the outgrowth of *L. murinus* is considered to impair gut metabolic function (34), therefore the depletion of *L. murinus* led by FMT may attenuate the microbial dysbiosis. Our metagenomic results are in line with the previously published studies that FMT could reshape the composition of both beneficial and harmful bacteria in the gut microbiome upon the anti-PD-1 treatment, which might result in the enhanced therapeutic efficacy.

Microbial gene functions and host metabolome were also reshaped by FMT in this study, which might benefit the efficacy of immunotherapy. Microbial gene pathways including nucleotides and amino acid biosynthesis pathways (e.g., pyrimidine deoxyribonucleotides, guanosine nucleotides, ornithine, isoleucine) were enriched after FMT, whereas methionine and SAM biosynthesis pathways were significantly downregulated **(Fig. 3A, B**). Methionine is involved in the pathogenesis of cancer (35), and negatively related to the efficacy of radiotherapy (36). SAM, a universal methyl donor, is formed from methionine and has been reported to be associated with metastasis and recurrence in colorectal cancer patients (37). Inhibition of the production of methionine and SAM might contribute to the tumor regression. Furthermore, our metabolomics analysis showed higher abundance of aspirin which can inhibit the growth of *Fusobacterium nucleatum* (a detrimental bacteria species which aggravates colorectal cancer) after FMT treatment (38). Likewise, punicic acid was regulated upon FMT. The potent anti-tumor effect of punicic acid might play a role in tumor control (39, 40). Lastly, the abundance of several amino acids was also decreased in the plasma, including glycine, serine, and cysteine (**Fig. 4C, E, Fig. S3**). Previous research reported that the growth and proliferation of cancer cells require serine and glycine, and limiting exogenous serine and glycine could inhibit tumor growth in mouse models of colon cancer (41, 42). To summarize, the enhanced efficacy of anti-PD-1 therapy led by FMT might be mediated by the altered microbial genome and blood metabolome.

The limitations of this study include the lack of experimental validation of the aforementioned bacterial species and metabolic pathways. Also, the synergistic effect exerted in mouse model may vary from that in the clinic. Further clinical investigation is being conducted in our laboratory and is anticipated to shed light on the detailed mechanisms of the promising combined use of FMT and anti-PD-1 therapy.

## 5. Conclusion

In summary, our study provides novel insight into the synergetic effects of microbiota transplantation and anti-PD-1 therapy in treating colorectal cancer, including the remodeling of gut microbiota and plasma metabolome. Out results suggest that *Bacteroides*, including the FMT-increased *B. thetaiotaomicron, B. fragilis*, and *B. cellulosilyticus* and decreased *B. ovatus* might play a role in improving the efficacy of anti-PD-1 therapy. This work provides a mechanistic basis to further understand the role of FMT combined with anti-PD-1 therapy in treating various cancer types including colorectal cancer.

## Acknowledgements

We thank Yan Kou, Xiaomin Xu, Bangzhuo Tong (Xbiome) and Wenting Liu (Sun Yat-Sen University) for their kind help with data analysis. This work is supported by National Key Research and Development Program of China (2020YFA0907803) and Xbiome Biotech Co. Ltd.

## Author contributions

HH, YT, YY and WZ conceived the study. JH, XZ, WK and HH conducted the experiments. JH, XZ, HH, WK, YM and HZ performed data analysis and interpretation. YC, YH, YT, WZ and YY supervised and financially supported the study. JH, XZ, WK, WZ and YY wrote the manuscript with extensive input from all authors.

## Data availability statement

Metagenomic raw data is available at https://www.ncbi.nlm.nih.gov/bioproject/PRJNA799796.

