## Supplementary figures for "Metagenomic and Metabolomic Analyses Reveal Synergistic Effects of Fecal Microbiota Transplantation and Anti-PD-1 Therapy on Treating Colorectal Cancer"

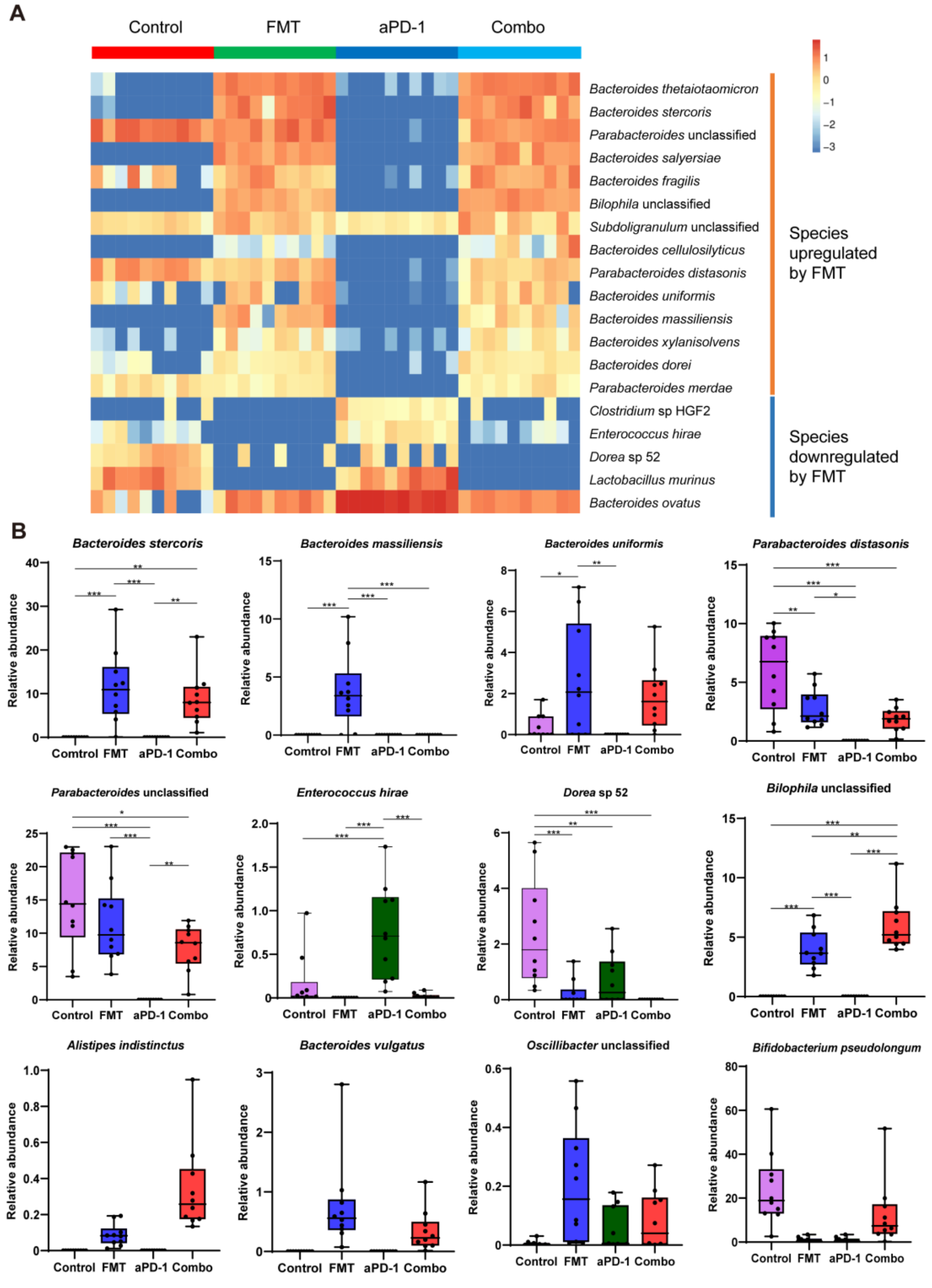


**Figure S1.** FMT altered the composition of gut microbiota in CT-26 tumor-bearing mice receiving anti-PD-1 therapy. (A) Heatmap of differentially abundant bacterial species. (B) Abundance of specific bacterial species in different groups. Data are represented as mean ± SD. *, p<0.05; **, p<0.01; ***, p<0.001.

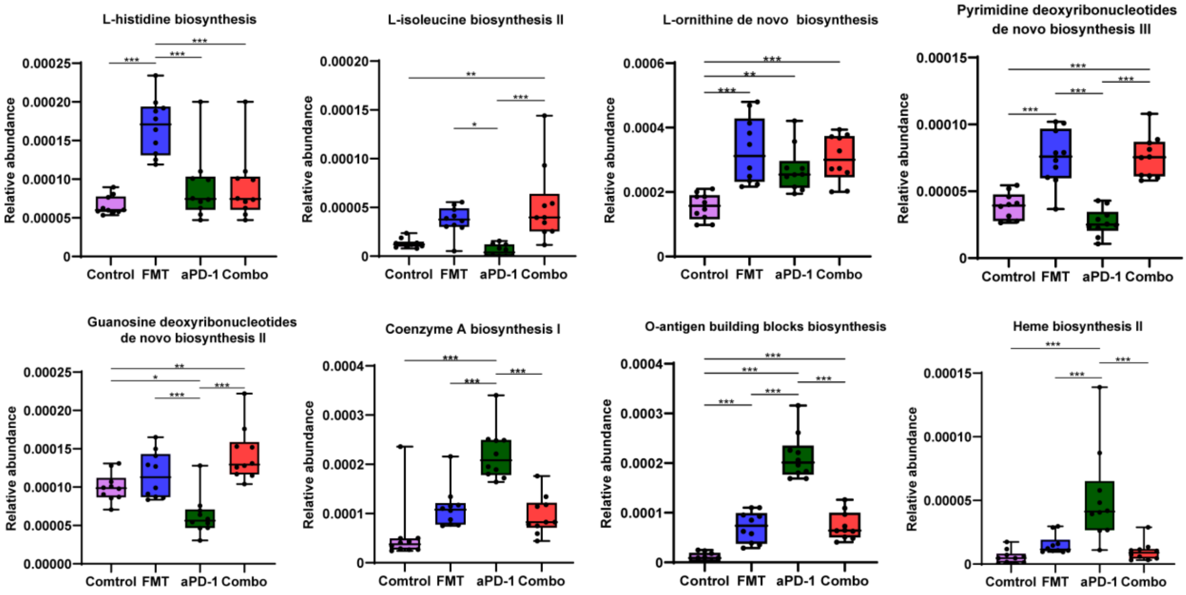


**Figure S2.** Abundance of significant gene pathways following different treatments. *, p<0.05; **, p<0.01; ***, p<0.001.

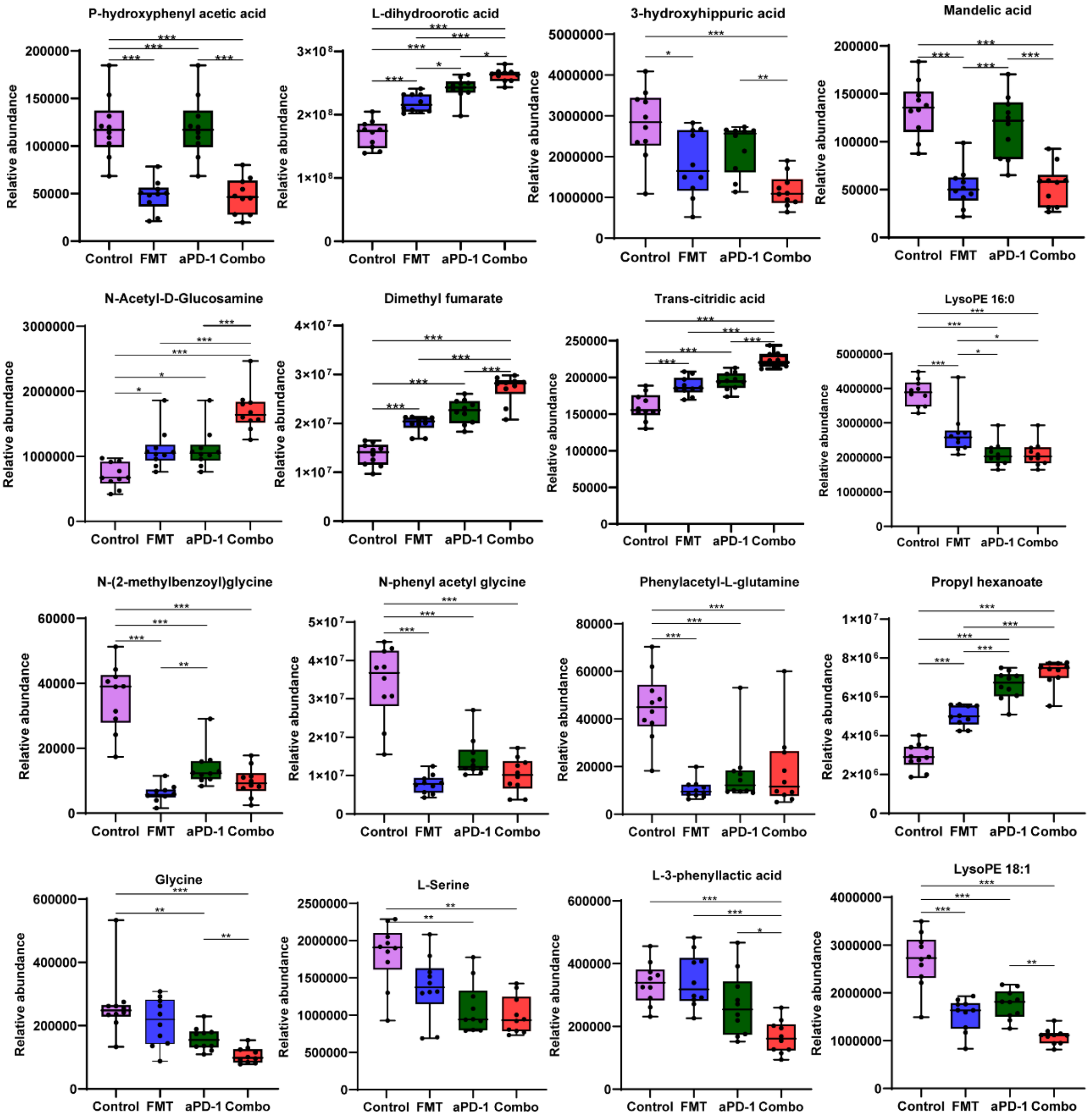


**Figure S3**. Abundance of significant metabolites following different treatments. Data are represented as mean ± SD. *, $p<0.05$; **, $p<0.01$; ***, $p<0.001$.
